# Assessing 16S marker gene survey data analysis methods using mixtures of human stool sample DNA extracts

**DOI:** 10.1101/400226

**Authors:** Nathan D Olson, M. Senthil Kumar, Shan Li, Stephanie Hao, Winston Timp, Marc L. Salit, O.Colin Stine, Hector Corrada Bravo

**Affiliations:** Biosystems and Biomaterials Division, National Institute of Standards and Technology, 100 Bureau Dr., Gaithersburg, Maryland, 20899 USA.; Center for Bioinformatics and Computational Biology, University of Maryland, College Park, 8314 Paint Branch Dr. College Park, Maryland, 20742 USA.; University of Maryland Institute of Advanced Computer Studies, University of Maryland, College Park, 8223 Paint Branch Dr. College Park, Maryland, 20742 USA.; Department of Epidemiology and Public Health, University of Maryland School of Medicine, 655 W. Baltimore Street, Baltimore, Maryland, 21201 USA.; Department of Biomedical Engineering, Johns Hopkins University, 720 Rutland Ave., Baltimore, Maryland, 21205 USA.; Joint Initiative for Metrology in Biology, National Institute of Standards and Technology, 443 Via Ortega, Stanford, CA, 94305 USA.; Department of Computer Science, University of Maryland, College Park, 8223 Paint Branch Dr. College Park, Maryland, 20742 USA.

**Keywords:** 16S rRNA gene, assessment, bioinformatic pipeline, normalization, differential abundance

## Abstract

**Background:** Analysis of 16S rRNA marker-gene surveys, used to characterize prokaryotic microbial communities, may be performed by numerous bioinformatic pipelines and downstream analysis methods. However, there is limited guidance on how to decide between methods, appropriate data sets and statistics for assessing these methods are needed. We developed a mixture dataset with real data complexity and an expected value for assessing 16S rRNA bioinformatic pipelines and downstream analysis methods. We generate an assessment dataset using a two-sample titration mixture design. The sequencing data were processed using multiple bioinformatic pipelines, i) DADA2 a sequence inference method, ii) Mothur a *de novo* clustering method, and iii) QIIME with open-reference clustering. The mixture dataset was used to qualitatively and quantitatively assess count tables generated using the pipelines.

**Results:** The qualitative assessment was used to evalute features only present in unmixed samples and titrations. The abundance of Mothur and QIIME features specific to unmixed samples and titrations were explained by sampling alone. However, for DADA2 over a third of the unmixed sample and titration specific feature abundance could not be explained by sampling alone. The quantitative assessment evaluated pipeline performance by comparing observed to expected relative and differential abundance values. Overall the observed relative abundance and differential abundance values were consistent with the expected values. Though outlier features were observed across all pipelines.

**Conclusions:** Using a novel mixture dataset and assessment methods we quantitatively and qualitatively evaluated count tables generated using three bioinformatic pipelines. The dataset and methods developed for this study will serve as a valuable community resource for assessing 16S rRNA marker-gene survey bioinformatic methods.

## Background

Targeted sequencing of the 16S rRNA gene, commonly known as 16S rRNA markergene-surveys, is a commonly used method for characterizing microbial communities, microbiomes. The 16S rRNA marker-gene-survey measurement process includes molecular (e.g. PCR and sequencing) and computational steps (e.g., sequence clustering) [1]. Molecular steps are used to selectively target and sequence the 16S rRNA gene from prokaryotic organisms within a sample. The computational steps convert the raw sequence data into a matrix with feature (e.g., operational taxonomic units) relative abundance values, feature abundance relative to all other features, for each sample [1]. Both molecular and computational measurement process steps contribute to the overall measurement bias and dispersion [2, 1, 3]. Proper measurement method evaluation allows for the characterization of how individual steps impact the measurement processes as a whole and determine where to focus efforts for improving the measurement process. Appropriate datasets and methods are needed to evaluate the 16S rRNA marker-gene-survey measurement process. A sample or dataset with “ground truth” is needed to characterize measurement process accuracy. Numerous studies have evaluated quantitative and qualitative characteristics of the 16S rRNA measurement process using mock communities, simulated data, and environmental samples.

To assess the qualitative characteristics of the 16S rRNA sequencing measurement process mock communities are commonly used [4]. As the number of organisms in the mock community is known, the total number of features can be compared to the expected value. The number of observed features in a mock community is significantly higher than the expected number of organism [5]. The higher than expected number of features is often attributed to sequencing and PCR artifacts as well as reagent contaminants [3, 6]. A notable exception to this is mock community benchmarking studies evaluating sequencing inference method, such as DADA2 [7]. Sequence inference methods aim to reduce the number of sequence artifact features. While mock communities have an expected number of features and composition, they lack the feature diversity and relative abundance dynamic range of real samples [4].

The quantitative characteristics of 16S rRNA sequence data are normally assessed using mock communities and simulated data. Mock communities of equimolar and staggered concentration are used to assess relative abundance estimate quantitative accuracy [5]. Results from relative abundance estimates using mock communities generated from mixtures of single organism DNA have shown taxonomic specific effects where individual taxa are under or over represented in a sample. For example Gram-negative bacteria have higher extraction efficiency compared to Grampositive bacteria [8, 9]. Mismatches in the primer binding sites are also responsible for taxonomic specific biases [3, 10, 11]. Additionally, taxon specific biases due to sequence template properties such as GC content, secondary structure, and gene flanking regions have been observed [12, 13, 11]. Simulated count tables have been used to assess differential abundance method (fold change differences in relative abundance), where specific taxa are artificially overrepresented in one set of samples compared to another [14]. Using simulated data to assess log fold-change estimates only evaluates computational steps of the measurement process.

Quantitative and qualitative assessment can also be performed using sequence data generated from mixtures of environmental samples. While simulated data and mock communities are useful in evaluating and benchmarking new methods one needs to consider that methods optimized for mock communities and simulated data are not necessarily optimized for the sequencing error profile and feature diversity of real samples. Data from environmental samples, which are real samples, are often used to benchmark new molecular laboratory and computational methods. However, without an expected value to compare to, only measurement precision, agreement with other methods, can be evaluated. By mixing environmental samples, expected values are calculated using information from the unmixed samples and mixture design. Mixtures of environmental samples were previously used to evaluate gene expression measurements [15, 16, 17].

In the present study, we developed a mixture dataset of extracted DNA from human stool samples for assessing 16S rRNA sequencing. The mixture datasets were processed using three bioinformatic pipelines. We developed metrics for qualitative and quantitative assessment of the bioinformatic pipeline results. The quantitative results were similar across pipelines, but the qualitative results varied by pipeline. We have made both the dataset and metrics developed in this study publicly available for evaluating bioinformatic pipelines.

## Results

### Two-Sample Titration Design

Samples collected at multiple timepoints during a Enterotoxigenic *E. coli* (ETEC) vaccine trial [18] were used to generate a two-sample titration dataset for assessing the 16S rRNA marker-gene survey measurement process. Samples from five trial participants were selected for our two-sample titration dataset. Trial participants (subjects) and sampling timepoints were selected based on *E. coli* abundance data collected using qPCR and 16S rRNA sequencing from Pop et al. [19]. Only individuals with no *E. coli* detected in samples collected from trial participants prior to ETEC exposure (PRE) were used for our two-samples titrations. Post ETEC exposure (POST) samples were identified as the timepoint after exposure to ETEC with the highest *E. coli* concentration for each subject (Fig. 1A). Due to limited sample availability, for E01JH0016 the timepoint with the second highest *E. coli* concentration was used as the POST sample. Independent titration series were generated for each subject, where POST samples were titrated into PRE samples with POST proportions of 1/2, 1/4, 1/8, 1/16, 1/32, 1/1,024, and 1/32,768 (Fig. 1B). Unmixed (PRE and POST) sample DNA concentration was measured using NanoDrop ND1000 (Thermo Fisher Scientific Inc. Waltham, MA USA). Unmixed samples were diluted to 12.5 *ng/µL* in tris-EDTA buffer before mixing.

**Figure 1.**
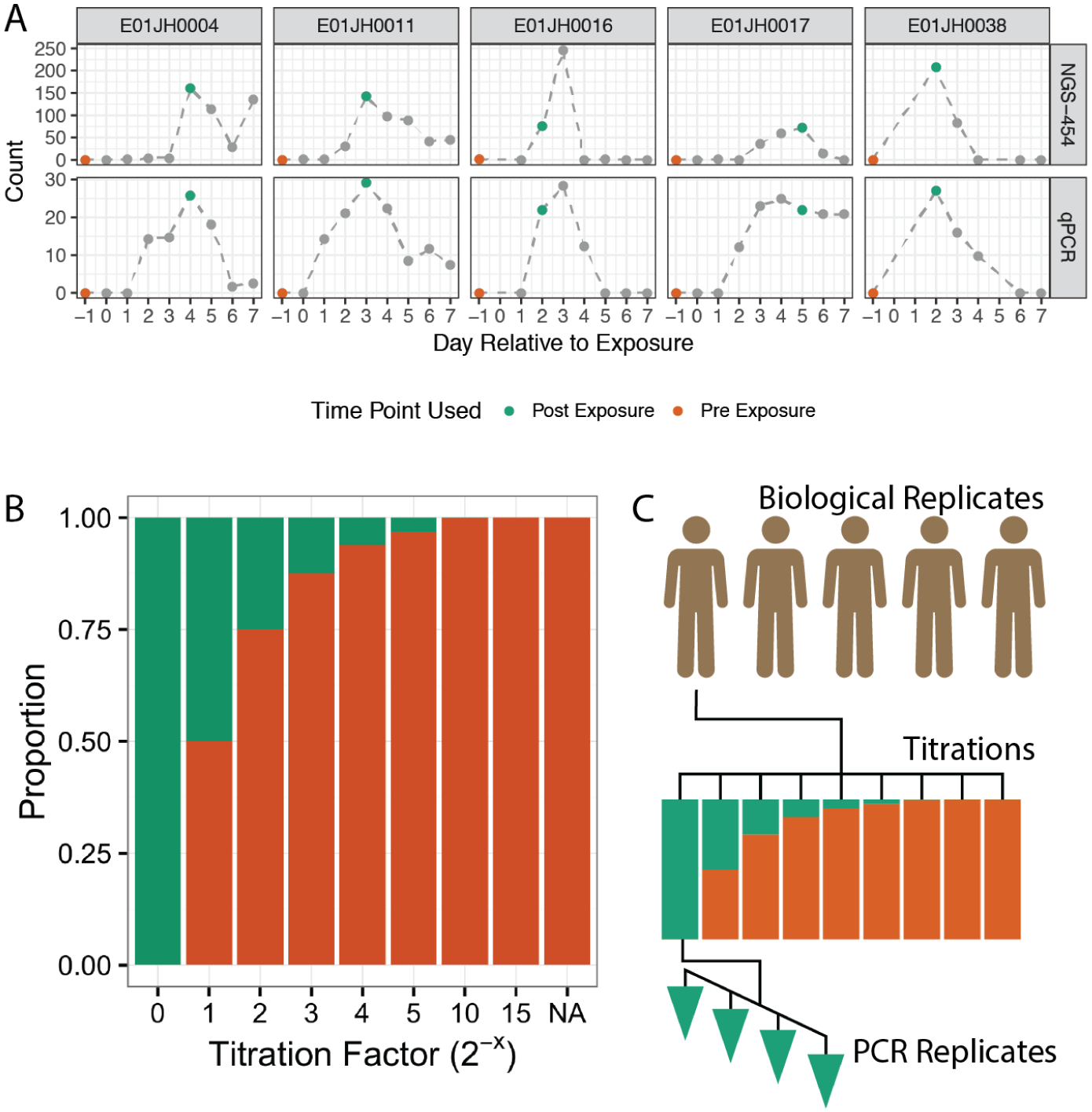
Sample selection and experimental design for the two-sample titration 16S rRNA marker-gene-survey assessment dataset. A) Preand post-exposure (PRE and POST) samples from five vaccine trial participants were selected based on *Escherichia coli* abundance measured using qPCR and 454 16S rRNA sequencing (454-NGS), data from Pop et al. [19]. Counts represent normalized relative abundance values for 454-NGS and copies of the heat-labile toxin gene per *µL*, a marker gene for ETEC, for qPCR. PRE and POST samples are indicated with orange and green data points, respectively. Grey points are other samples from the vaccine trial time series. B) Proportion of DNA from PRE and POST samples in titration series samples. PRE samples were titrated into POST samples following a *log*_2_ dilution series. The NA titration factor represents the unmixed PRE sample. C) PRE and POST samples from the five vaccine trial participants, subjects, were used to generate independent two-sample titration series. The result was a total of 45 samples, 7 titrations + 2 unmixed samples times 5 subjects. Four replicate PCRs were performed for each of the 45 samples resulting in 190 PCRs.

For our two-sample titration mixture design, the expected feature relative abundance can be calculated using equation (1), where *θ*_*i*_, is the proportion of POST DNA in titration *i*, *q*_*ij*_ is the relative abundance of feature *j* in titration *i*, and the relative abundance of feature *j* in the unmixed PRE and POST samples is *q*_*pre,j*_ and *q*_*post,j*_.

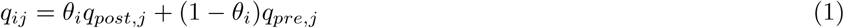

### Dataset characteristics

We first characterize the number of reads per sample and base quality score distribution. The number of reads per sample and distribution of base quality scores by position was consistent across subjects (Fig. 2). Two barcoded experimental samples had less than 35,000 reads. The rest of the samples with less than 35,000 reads were no template PCR controls (NTC). Excluding one failed reaction with 2,700 reads and NTCs, there were 8.9548 *×* 10^4^ (3195-152267) sequences per sample, median and range. Forward reads had consistently higher base quality scores relative to the reverse reads with a narrow overlap region with high base quality scores for both forward and reverse reads (Fig. 2B).

**Figure 2.**
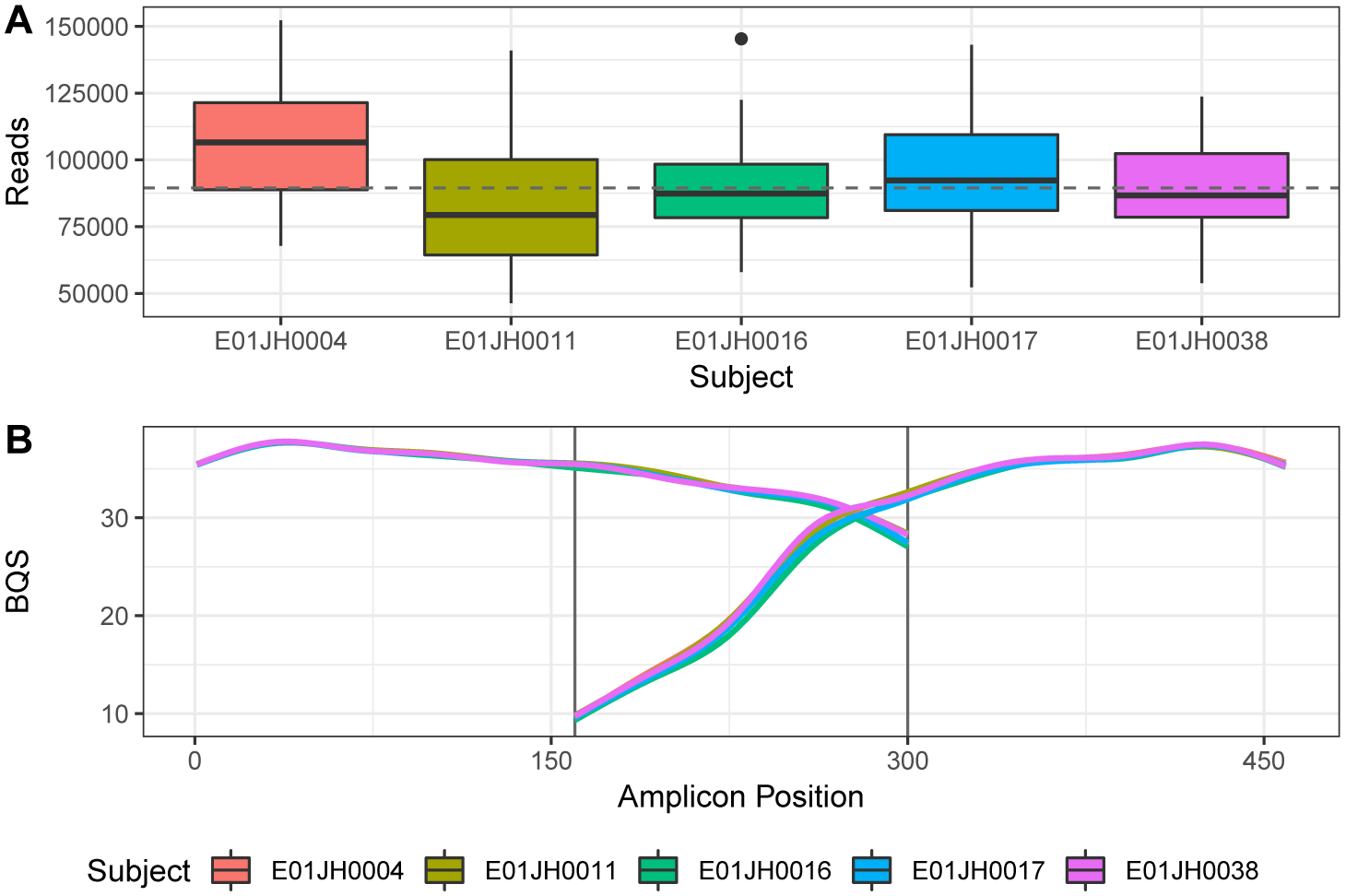
Sequence dataset characteristics. (A) Distribution in the number of reads per barcoded sample (Library Size) by individual. Boxplots summarize data distribution with horizontal bar as median, boxes indicating interquartile range, whiskers *±*1.5*× IQR*, and black points outliers. The dashed horizontal line indicates overall median library size. Excluding one PCR replicate from subject E01JH0016 titration 5 that had only 3,195 reads. (B) Smoothing spline of the base quality score (BQS) across the amplicon by subject. Vertical lines indicate approximate overlap region between forward and reverse reads. Forward reads go from position 0 to 300 and reverse reads from 464 to 164.

The resulting count tables generated using the four bioinformatic pipelines were characterized for number of features, sparsity, and filter rate (Table 1, Figs. 3B). The pipelines evaluated employ different approaches for handling low quality reads resulting in large differences in drop-out rate and the fraction of raw sequences not included in the count table (Table 1). QIIME pipeline has the highest drop-out rate and number of features per sample but fewer total features than Mothur. The targeted amplicon region has a relatively small overlap region, 136 bp for 300 bp paired-end reads, compared to other commonly used amplicons [20, 21]. The high drop-out rate is due to low basecall accuracy at the ends of the reads especially the reverse reads resulting in a high proportion of unsuccessfully merged reads pairs (Fig. 2B). Furthermore, increasing the drop-out rate, QIIME excludes singletons, features only observed once in the dataset, to remove potential sequencing artifacts from the dataset. QIIME and DADA2 pipelines were similarly sparse (the fraction of zero values in count tables) despite differences in the number of features and dropout rate. The expectation is that this mixture dataset will be less sparse relative to other datasets. This is due to the redundant nature of the samples where the 35 titration samples are derived directly from the 10 unmixed samples, along with four PCR replicates for each sample. With sparsity greater than 0.9 for the three pipelines it is unlikely that any of the pipelines successfully filtered out a majority of the sequencing artifacts.

**Figure 3.**
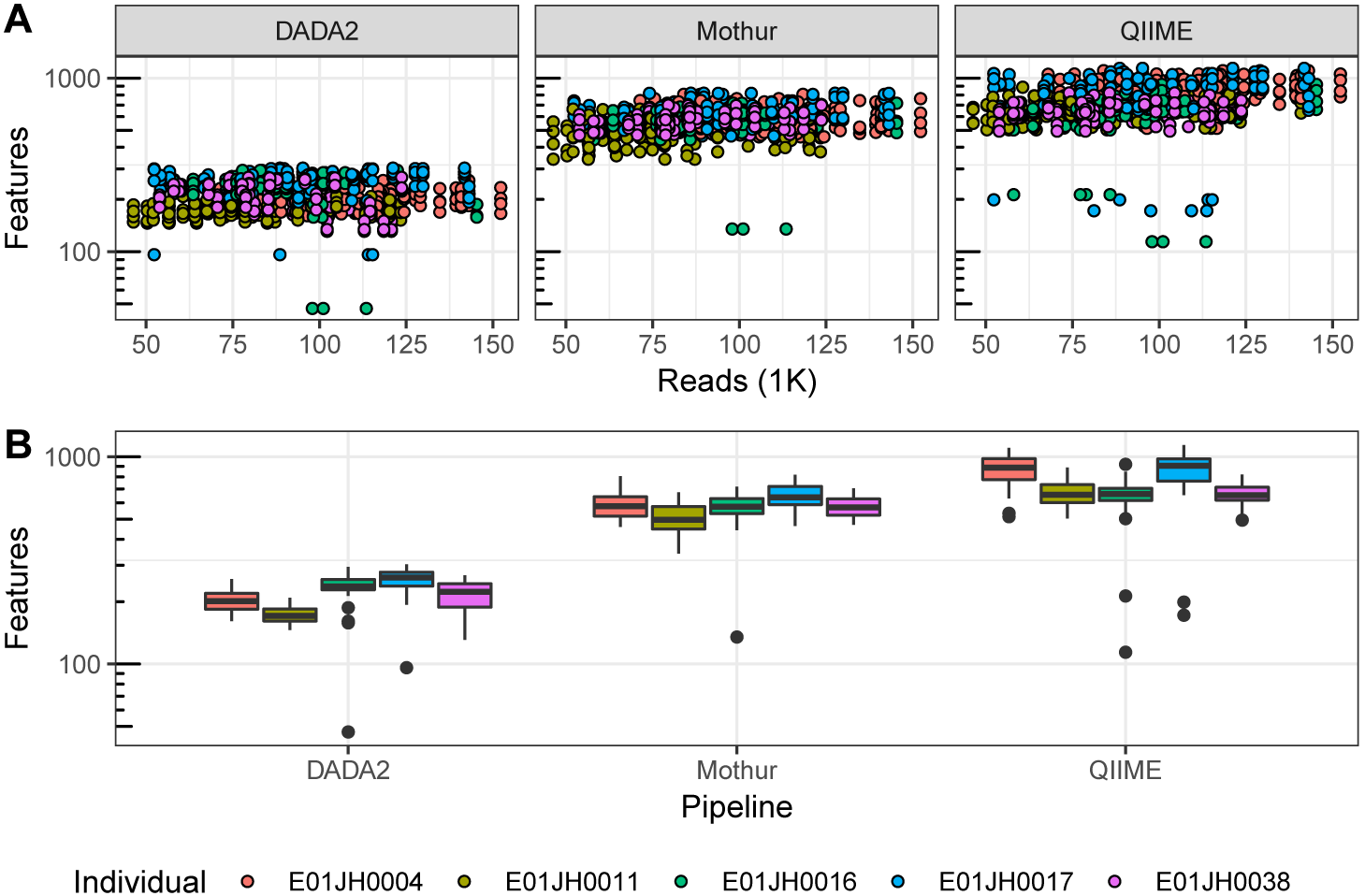
Relationship between the number of reads and features per sample by bioinformatic pipeline. (A) Scatter plot of observed features versus the number of reads per sample. (B) Observed feature distribution by pipeline and individual. Excluding one PCR replicate from subject E01JH0016 titration 5 with only 3,195 reads, and the Mothur E01JH0017 titration 4 (all four PCR replicates), with 1,777 observed features.

**Table 1.**
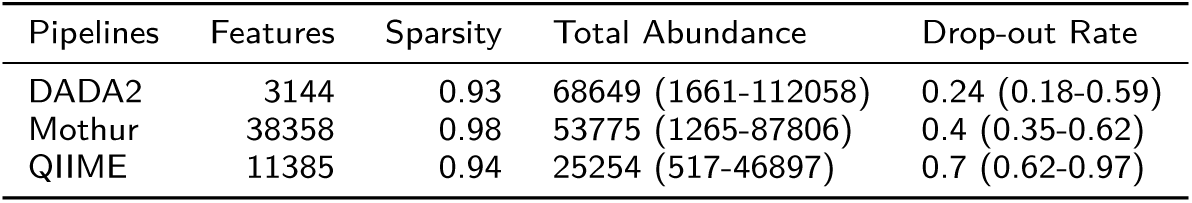
Summary statistics for the different bioinformatic pipelines. DADA2 is a denoising sequence inference pipeline, QIIME is an open-reference clustering pipeline, and Mothur is a de-novo clustering pipeline. No template controls were excluded from summary statistics. Sparsity is the proportion of 0’s in the count table. Features is the total number of OTUs (QIIME and Mothur) or SVs (DADA2) in the count. Sample coverage is the median and range (minimum-maximum) per sample total abundance. Drop-out rate is the proportion of reads removed while processing the sequencing data for each bioinformatic pipeline.

Dataset taxonomic assignments also varied by pipeline (Fig. 4). Phylum and order relative abundance is similar across pipelines (Fig. 4A & B). The observed differences are attributed to different taxonomic classification methods and databases used by the pipelines. DADA2 and QIIME pipelines differed from Mothur and QIIME for Proteobacteria and Bacteriodetes. Regardless of the relative abundance threshold, for genus sets most genera were unique to individual pipelines (Fig. 4C & D). Sets, shared taxa between pipelines, with QIIME had the fewest genera, excluding the DADA2-QIIME set. QIIME was the only pipeline to use open-reference clustering and the Greengenes database. Mothur and DADA2 both used the SILVA dataset. The Mothur and DADA2 pipeline use different implmentations of the RDP naÏve Bayesian classifier, which may be partially responsible for the Mothur, unclustered, and DADA2 differences.

**Figure 4.**
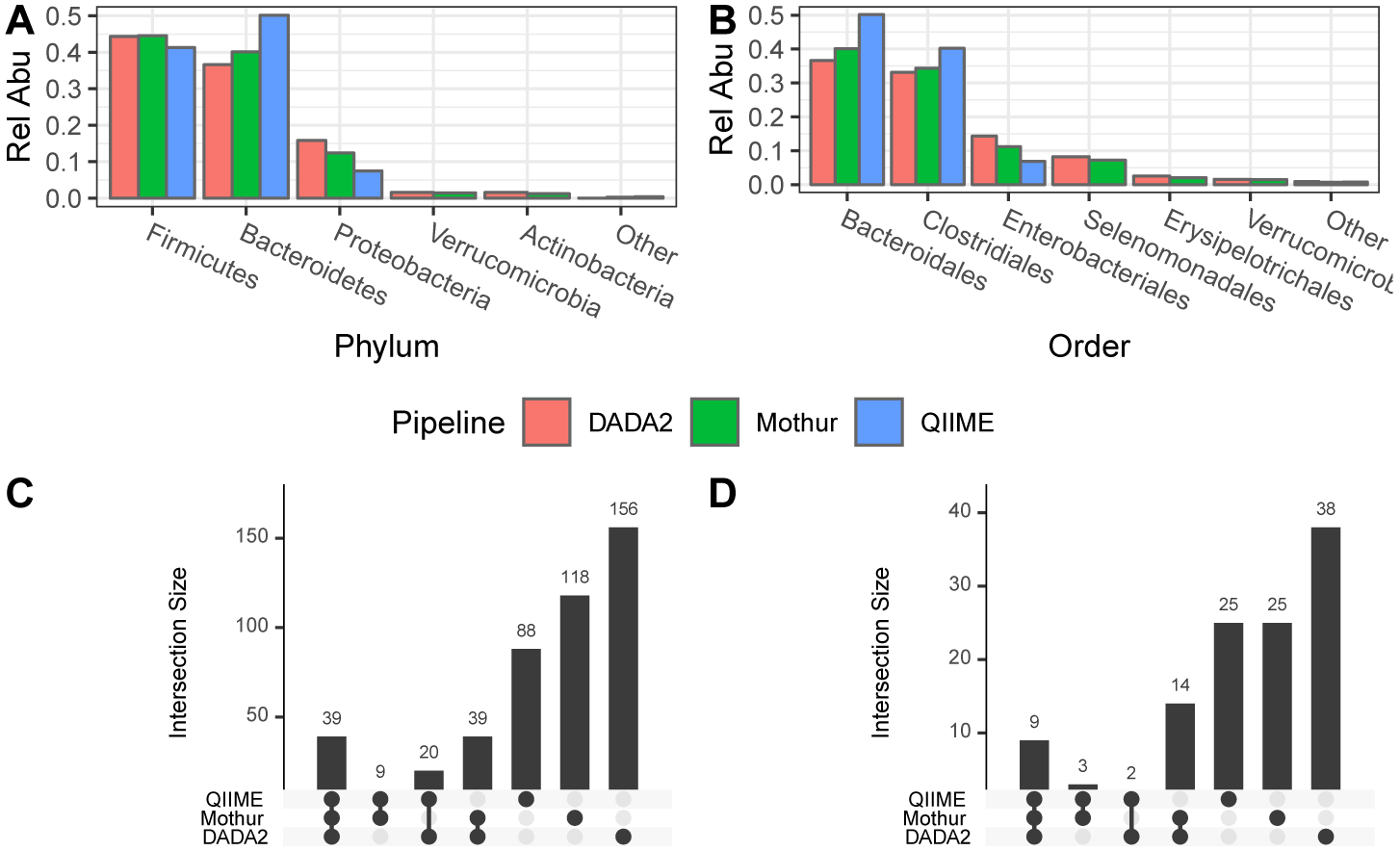
Comparison of dataset taxonomic composition across pipelines. Phylum (A) and Order (B) relative abundance by pipeline. Taxonomic groups with less than 1% total relative abundance were grouped together and indicated as other. Pipeline genus-level taxonomic assignment set overlap for the all features (C) and the upper quartile genera by relative abundance for each pipeline (D).

### Titration Series Validation

To validate the two-sample titration dataset for use in abundance assessment we evaluated two assumptions about the titrations: 1. The samples were mixed volumetrically in a *log*_2_ dilution series according to the mixture design. 2. The unmixed PRE and POST samples have the same proportion of prokaryotic DNA. The stool samples used to generate the mixtures have both eukaryotic (primarily human) DNA and prokaryotic DNA. If the proportion of prokaryotic DNA differs between the unmixed samples, then the amount of DNA from the unmixed samples in a titration targeted by 16S rRNA gene sequencing is not consistent with the mixture design. To validate the sample volumetric mixing exogenous DNA was spiked into the unmixed samples before mixing and quantified using qPCR. To evaluate if the PRE and POST samples had the same proportion of prokaryotic DNA total prokaryotic DNA in the titrations samples was quantified using a qPCR assay targeting the 16S rRNA gene.

#### Spike-in qPCR results

Titration series volumetric mixing was validated by quantifying ERCC plasmids spiked into the POST samples using qPCR. The qPCR assay standard curves had a high level of precision with *R*^2^ values close to 1 and amplification efficiencies between 0.84 and 0.9 for all standard curves indicating the assays were suitable for validating the titration series volumetric mixing (Table 2). For our *log*_2_ two-sample-titration mixture design the expected slope of the regression line between titration factor and Ct is 1, corresponding to a doubling in template DNA every PCR cycle. The qPCR assays targeting the ERCCs spiked into the POST samples had *R*^2^ values and slope estimates close to 1 (Table 2). Slope estimates less than one were attributed to assay standard curve efficiency less than 1 (Table 2). ERCCs spiked into PRE samples were not used to validate volumetric mixing as PRE sample proportion differences were too small for qPCR quantification. The expected *C*_*t*_ difference for the entire range of PRE concentrations in only 1. When considering the quantitative limitations of the qPCR assay these results confirm that the unmixed samples were volumetrically mixed according to the two-sample titration mixture design.

**Table 2.** ERCC Spike-in qPCR assay information and summary statistics. ERCC is the ERCC identifier for the ERCC spike-in, Assay is TaqMan assay, and Length and GC are the size and GC content of the qPCR amplicon. The Std. *R*^2^ and Efficiency (E) statistics were computed for the standard curves. *R*^2^ and slope for titration qPCR results for the titration series.

| Subject | ERCC | Assay | Length | Std. $R^2$ | E | $R^2$ | Slope |
| --- | --- | --- | --- | --- | --- | --- | --- |
| E01JH0004 | 012 | Ac03459877-a1 | 77 | 0.9996 | 86.19 | 0.98 | 0.92 |
| E01JH0011 | 157 | Ac03459958-a1 | 71 | 0.9995 | 87.46 | 0.95 | 0.90 |
| E01JH0016 | 108 | Ac03460028-a1 | 74 | 0.9991 | 87.33 | 0.95 | 0.84 |
| E01JH0017 | 002 | Ac03459872-a1 | 69 | 0.9968 | 85.80 | 0.89 | 0.93 |
| E01JH0038 | 035 | Ac03459892-a1 | 65 | 0.9984 | 86.69 | 0.95 | 0.94 |

#### Prokaryotic DNA Concentration

Observed changes in prokaryotic DNA concentration across titrations indicate the proportion of prokaryotic DNA from the unmixed PRE and POST samples in a titration is inconsistent with the mixture design (Fig. 5). A qPCR assay targeting the 16S rRNA gene was used to quantify the concentration of prokaryotic DNA in the titrations. An in-house standard curve with concentrations of 20 ng/ul, 2ng/ul, and 0.2 ng/ul was used, with efficiency 91.49, and *R*^2^ 0.999. If the proportion of prokaryotic DNA is the same between PRE and POST samples the slope of the concentration estimates across the two-sample titration would be 0. For subjects where the proportion of prokaryotic DNA is higher in the PRE samples, the slope will be negative, and positive when the proportion is higher for POST samples. The slope estimates are significantly different from 0 for all subjects excluding E01JH0011 (Fig. 5). These results indicate that the proportion of prokaryotic DNA is lower in POST when compared to the PRE samples for E01JH0004 and E01JH0017 and higher for E01JH0016 and E01JH0038.

**Figure 5.**
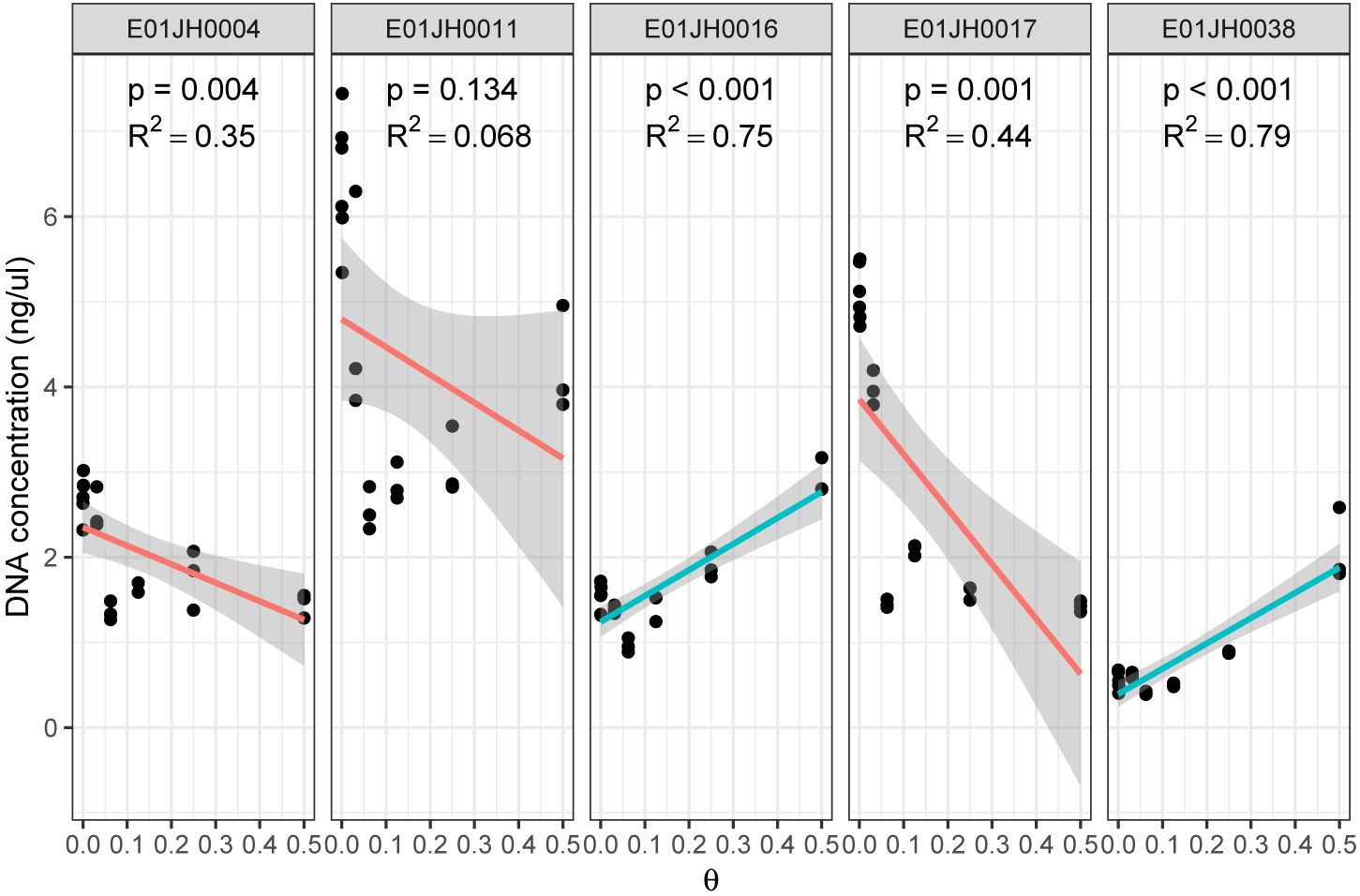
Prokaryotic DNA concentration (ng/ul) across titrations measured using a 16S rRNA qPCR assay. Separate linear models, Prokaryotic DNA concentration versus *θ* were fit for each individual, and *R*^2^ and p-values were reported. Red lines indicate negative slope estimates and blue lines positive slope estimates. p-value indicates significant difference from the expected slope of 0. The grey regions indicate the linear model 95% confidence interval. Multiple test correction was performed using the Benjamini-Hochberg method. One of the E01JH0004 PCR replicates for titration 3 (*θ* = 0.125) was identified as an outlier, with a concentration of 0.003, and was excluded from the linear model. The linear model slope was still significantly different from 0 when the outlier was included.

#### Theta Estimates

Human stool sample DNA extracts vary in the proportion of eukaryotic (primarily human) and prokaryotic DNA in the sample. To account for differences in the proportion of prokaryotic DNA in PRE and POST samples (Fig. 5) we inferred the proportion of POST sample prokaryotic DNA in a titration, *θ*, using the 16S rRNA sequencing data (Fig. 6). Overall the relationship between the inferred and mixture design *θ* values were consistent across pipelines but not subject whereas the *θ* estimate 95% CI varied by both subject and pipeline. For study subjects E01JH0004, E01JH0011, and E01JH0016 the inferred and mixture design *θ* values were in agreement, in contrast to study subjects E01JH0017 and E01JH0038. For E01JH0017 the inferred values were consistently less than the mixture design values. Whereas for E01JH0038 the inferred values were consistently greater than the mixture design values. These results were consistent with the qPCR prokaryotic DNA concentration results with significantly positive slopes for E01JH0004 and E01JH0016 and significantly negative slope for E01JH0038 (Fig. 5).

**Figure 6.**
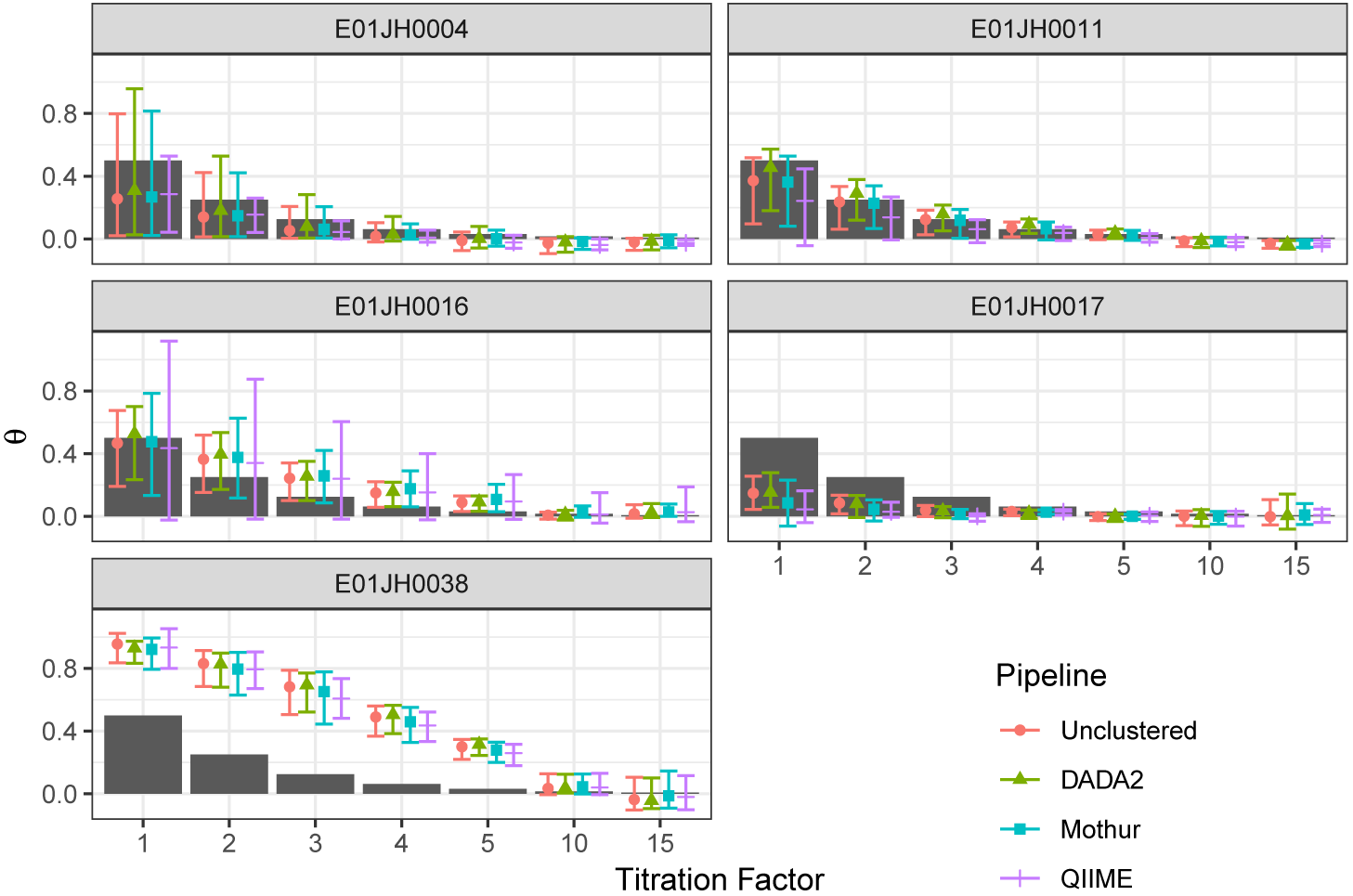
Theta estimates by titration, biological replicate, and bioinformatic pipeline. The points indicates mean estimate of 1000 bootstrap theta estimates and errorbars 95% confidence interval. The black bar indicate expected theta values. Theta estimates below the expected theta indicate that the titrations contain less than expected bacterial DNA from the POST sample. Theta estimates greater than the expected theta indicate the titration contains more bacterial DNA from the PRE sample than expected.

### Measurement Assessment

Next, we assessed the qualitative and quantitative nature of 16S rRNA measurement process using our two-sample titration dataset. For the qualitative assessment, we analyzed the relative abundance of features only observed in the unmixed samples or titrations. These features are not expected given the titration experimental design. The quantitative assessment evaluated relative and differential abundance estimates.

#### Qualitative Assessment

Unmixed- and titration-specific features were observed for all pipelines (titration-specific: Fig. 7A, unmixed-specific: Fig. 7B). For mixture datasets low abundance features present only in unmixed samples and mixtures are expected due to random sampling. For our two-sample titration dataset there were unmixed-specific features with expected counts not explained by sampling alone for all individuals and bioinformatic pipelines (Fig. 7C). However, the proportion of unmixed-specific features that could not be explained by sampling alone varied by bioinformatic pipeline. DADA2 had the highest proportion of unmixed-specific features not explained by sampling whereas QIIME had the lowest proportion. Consistent with the distribution of observed counts for titration-specific features more of the DADA2 features could not be explained by sampling alone compared to the other pipelines (Fig. 7D). Overall, the DADA2 count table had the largest number of observed features inconsistent with the titration experiment design, while the same phenomenon is significantly reduced in the other pipelines.

**Figure 7.**
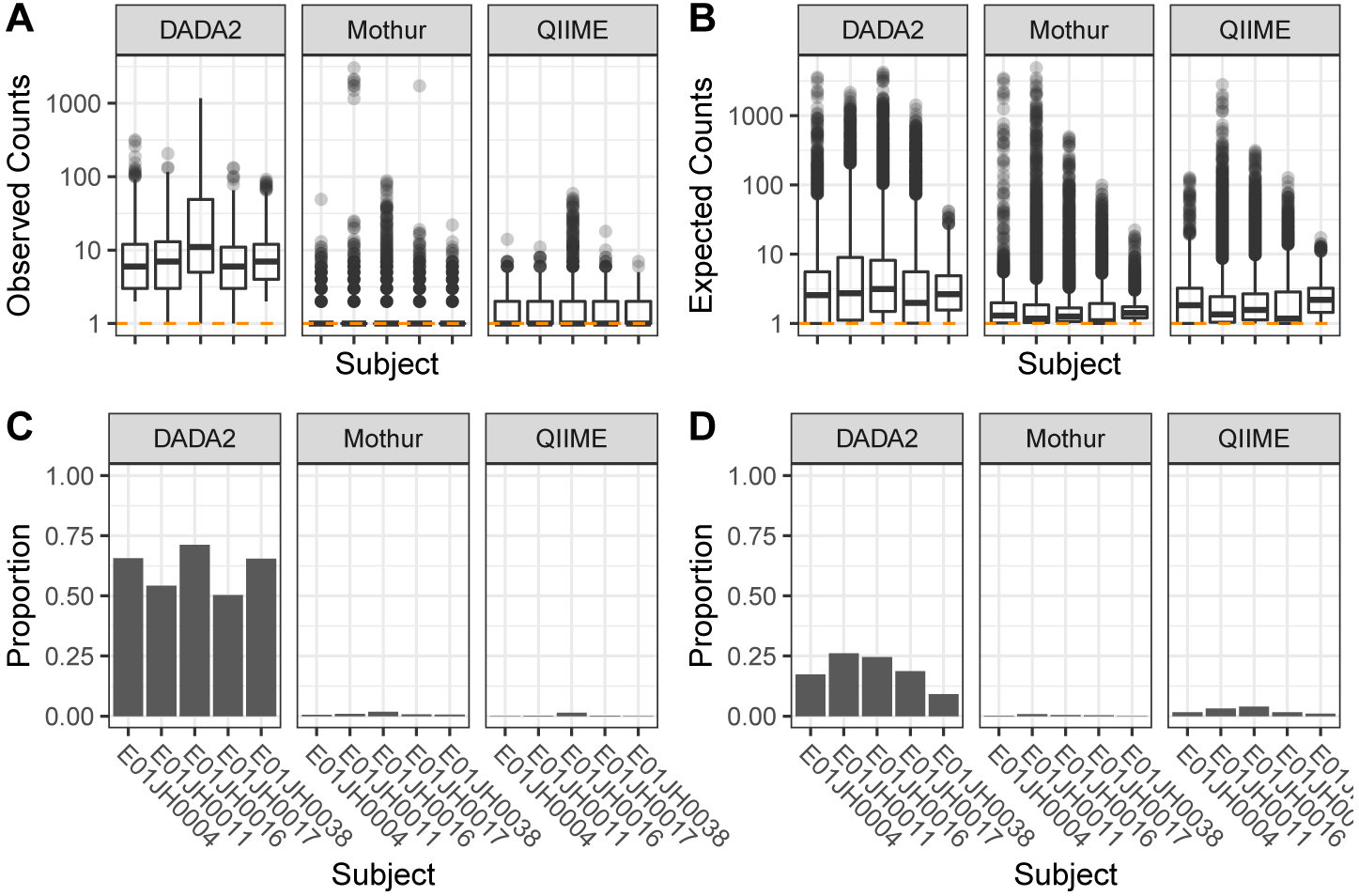
Distribution of (A) observed count values for titration-specific features and (B) expected count values for unmixed-specific features by pipeline and individual. The orange horizontal dashed line indicates a count value of 1. (C) Proportion of unmix-specific features and (D) titration-specific features with an adjusted p-value *<* 0.05 for the Bayesian hypothesis test and binomial test respectively. We failed to accept the null hypothesis when the p-value *<* 0.05, indicating that the discrepancy between the feature only being observed in the titrations or unmixed samples cannot be explained by sampling alone.

#### Quantitative Assessment

For the relative abundance assessment, we evaluated the consistency of the observed and expected relative abundance estimates for a feature and titration as well as feature-level bias and variance. The PRE and POST estimated relative abundance and inferred *θ* values were used to calculate titration and relative abundance error rates. Relative abundance error rate is defined as *|exp – obs|/exp*, where *exp* and *obs* is the expected and observed relative abundance. To control for biases in feature inference, the three pipelines were compared to an unclustered dataset. The unclustered count table was generated using the 40,000 most abundant features from Mothur’s initial preprocessing (see Methods for details). Unclustered pipeline *θ* estimates were used to calculate the error rates for all pipelines to prevent overfitting. Only features observed in all PRE and POST PCR replicates and PRE and POST specific features were included in the analysis (Table 3). PRE and POST specific features were defined as present in all four of the PRE or POST PCR replicates, respectively, but none of the PCR replicates for the other unmixed samples. There is lower confidence in PRE or POST feature relative abundance when the feature is not observed all 4 PCR replicates, therefore these features were not included in the analysis. Overall, agreement between inferred and observed relative abundance was high for all individuals and bioinformatic pipelines (Fig. 8A). The error rate distribution was similarly consistent across pipelines, including long tails (Fig. 8B)

**Figure 8.**
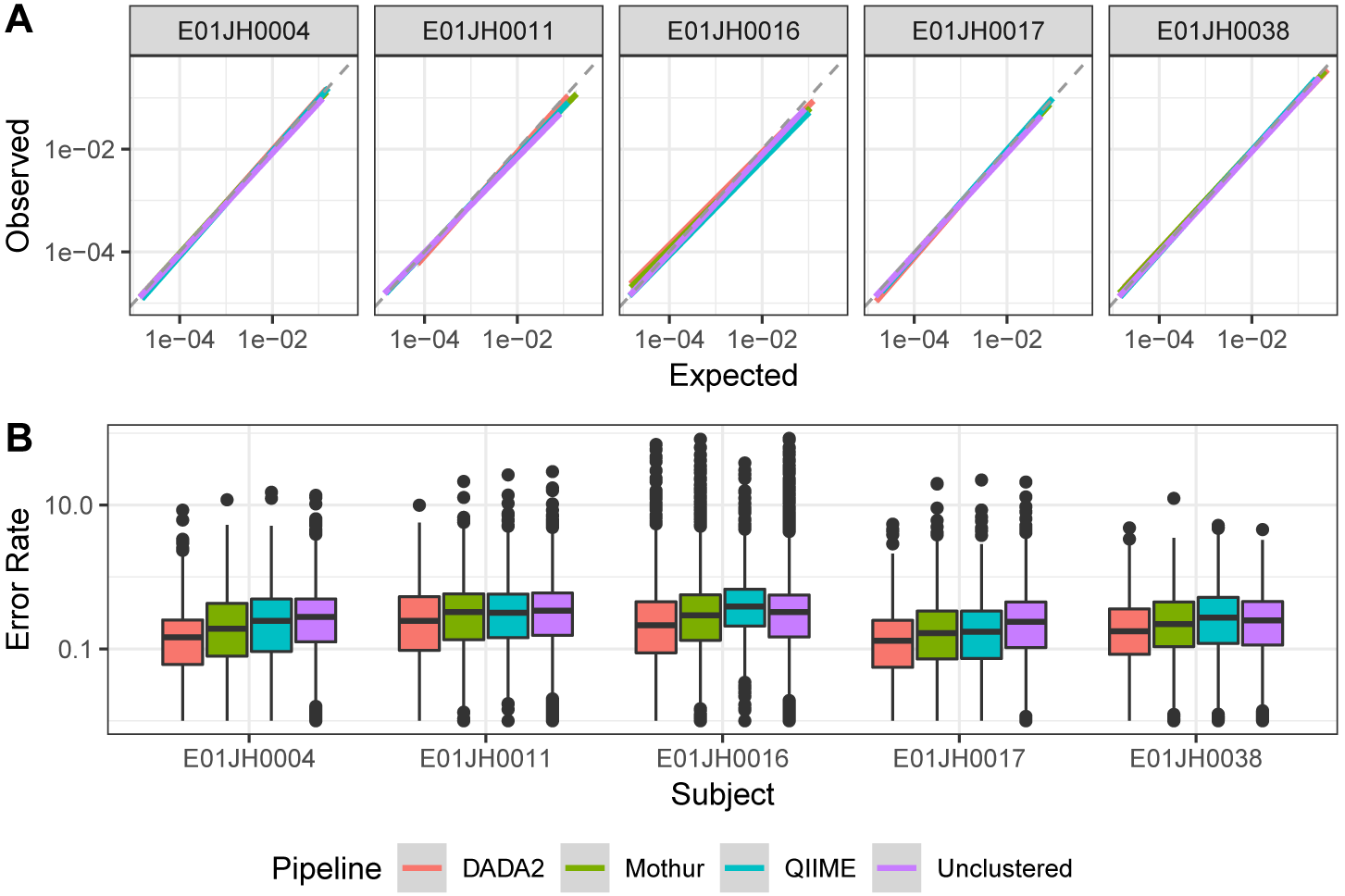
Relative abundance assessment. (A) A linear model of the relationship between the expected and observed relative abundance. The dashed grey line indicates expected 1-to-1 relationship. The plot is split by individual and color is used to indicate the different bioinformatic pipelines. A negative binomial model was used to calculate an average relative abundance estimate across the four PCR replicates. Points with observed and expected relative abundance values less than 1/median library size were excluded from the data used to fit the linear model. (B) Relative abundance error rate distribution by individual and pipeline.

**Table 3.**
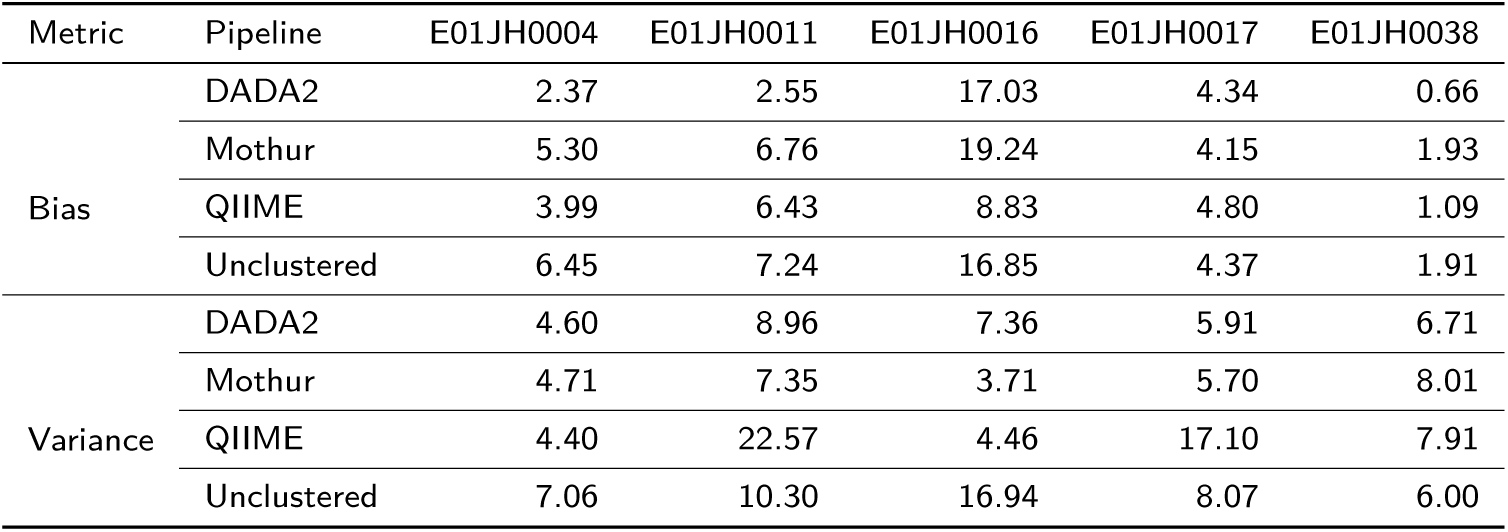
Maximum feature-level error rate bias (median error rate) and variance (robust COV) by pipeline and individual.

To assess quantitative accuracy, we compared the feature-level relative abundance error rate bias (median error rate, Fig. 9A) and variance (*RCOV* = (*IQR*)*/|median|* Fig. 9B) across pipelines and individuals using mixed effects models. Large bias and variance metric values were observed for all pipelines (Table 3). Features with large bias and variance metrics, 1.5 *× IQR* from the median, were deemed outliers. To prevent these outlier features from biasing the comparison they were not used to fit the mixed effects model. Multiple comparisons test (Tukey) was used to test for significant differences in feature-level bias and variance between pipelines. A one-sided alternative hypothesis was used to determine which pipelines had smaller feature-level error rate. The Mothur, DADA2, and QIIME feature-level bias were all significantly different from each other (*p <* 1 *×* 10^*-*8^). DADA2 had the lowest mean feature-level bias (0.2), followed by Mothur (0.28), with QIIME having the highest bias (0.33) (9B). Large variance metric values were observed for all individuals and pipelines (Table 3). The feature-level variance was not significantly different between pipelines, Mothur = 0.83, QIIME = 0.71 and DADA2 = 1 (Fig. 9B). We evaluated whether poor feature-level relative abundance metrics can be attributed to specific taxonomic groups or phylogenetic clades. While a significant overall phylogenetic signal was detected for both the bias and variance metric, no specific taxonomic groups or phylogenetic clades were identified with exceptionally poor performance in our assessment.

**Figure 9.**
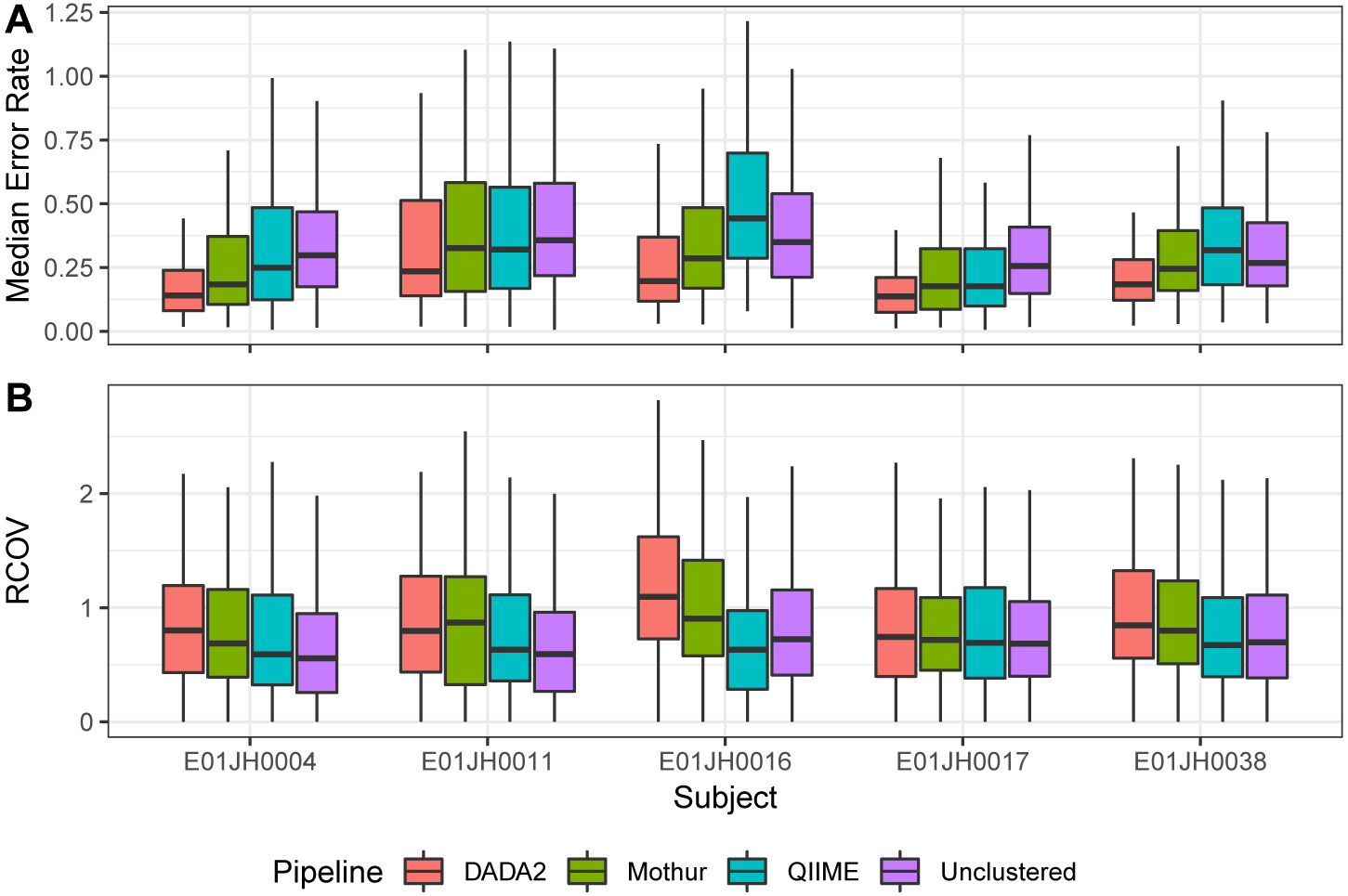
Comparison of pipeline relative abundance assessment feature-level error metrics. Distribution of feature-level relative abundance (A) bias metric median error rate and (B) variance robust coefficient of variation (*RCOV* = (*IQR*)*/ |median |*) by individual and pipeline. Boxplot outliers, 1.5*× IQR* from the median were excluded from the figure to prevent extreme metric values from obscuring metric value visual comparisons.

The agreement between log-fold change estimates and expected values were individual specific and consistent across pipelines (Fig. 10A). The individual specific effect was attributed to the fact that unlike relative abundance assessment the inferred *θ* values were not used to calculate expected values. Inferred *θ* values were not used to calculate the expected values because all of the titrations and the *θ* estimates for the higher titrations were included and they were not monotonically decreasing and therefore resulted in unrealistic expected log fold-change values, e.g., negative log-fold changes for PRE specific features. The log-fold change estimates and expected values were consistent across pipelines with one notable exception. For E01JH0011 the Mothur log fold-change estimates were more consistent with expected values than the other pipelines. However, as *θ* was not corrected for differences in the proportion of prokaryotic DNA between the unmixed PRE and POST samples it cannot be said whether Mothur’s performance was better than the other pipelines.

**Figure 10.**
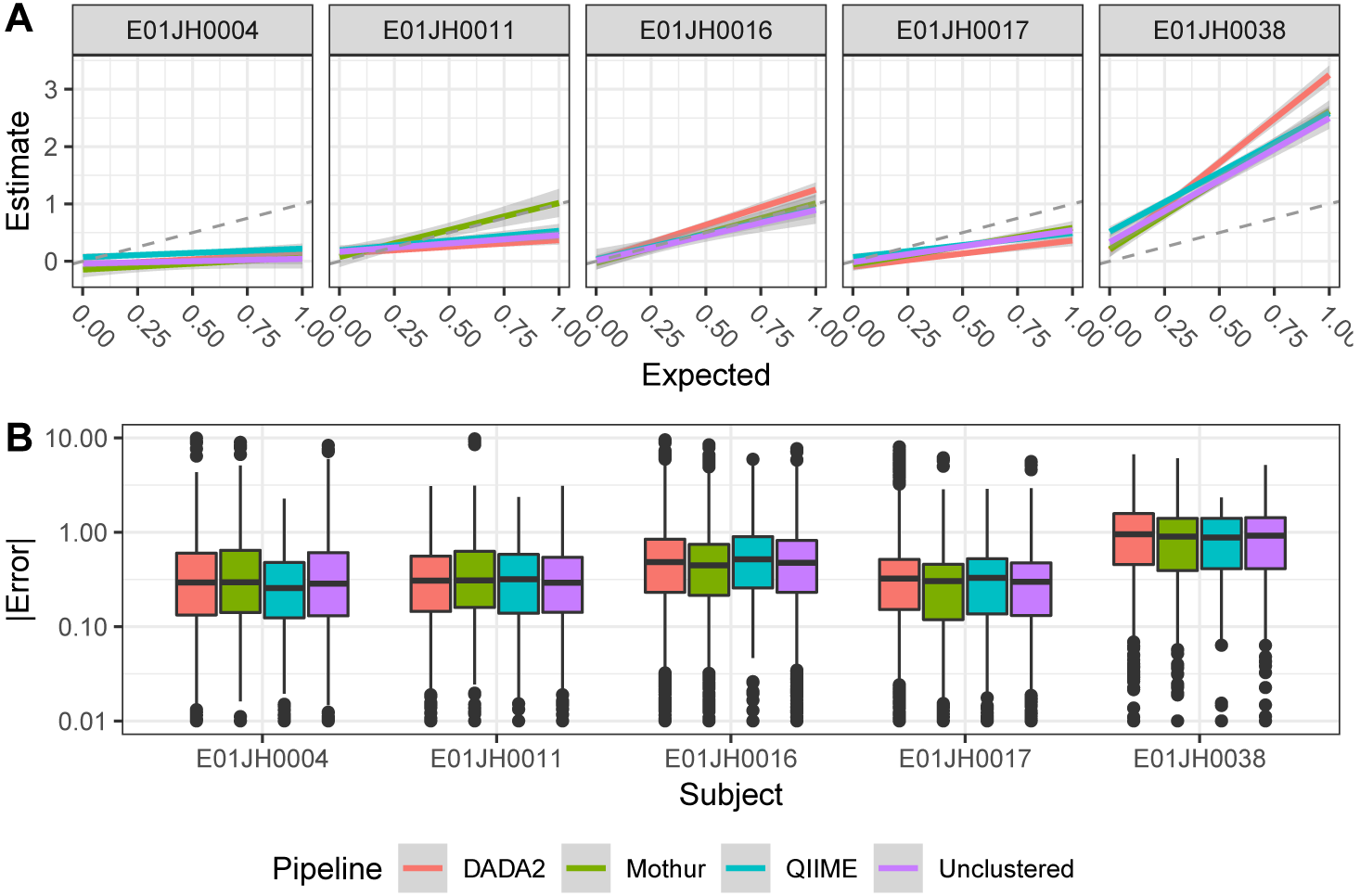
(A) Linear model or the relationship between log fold-change estimates and expected values for PRE-specific and PRE-dominant features by pipeline and individual, line color indicates pipelines. Dashed grey line indicates expected 1-to-1 relationship between the estimated and expected log fold-change. (B) Log fold-change error (*|*exp-est *|*) distribution by pipeline and individual.

**Figure 11.**
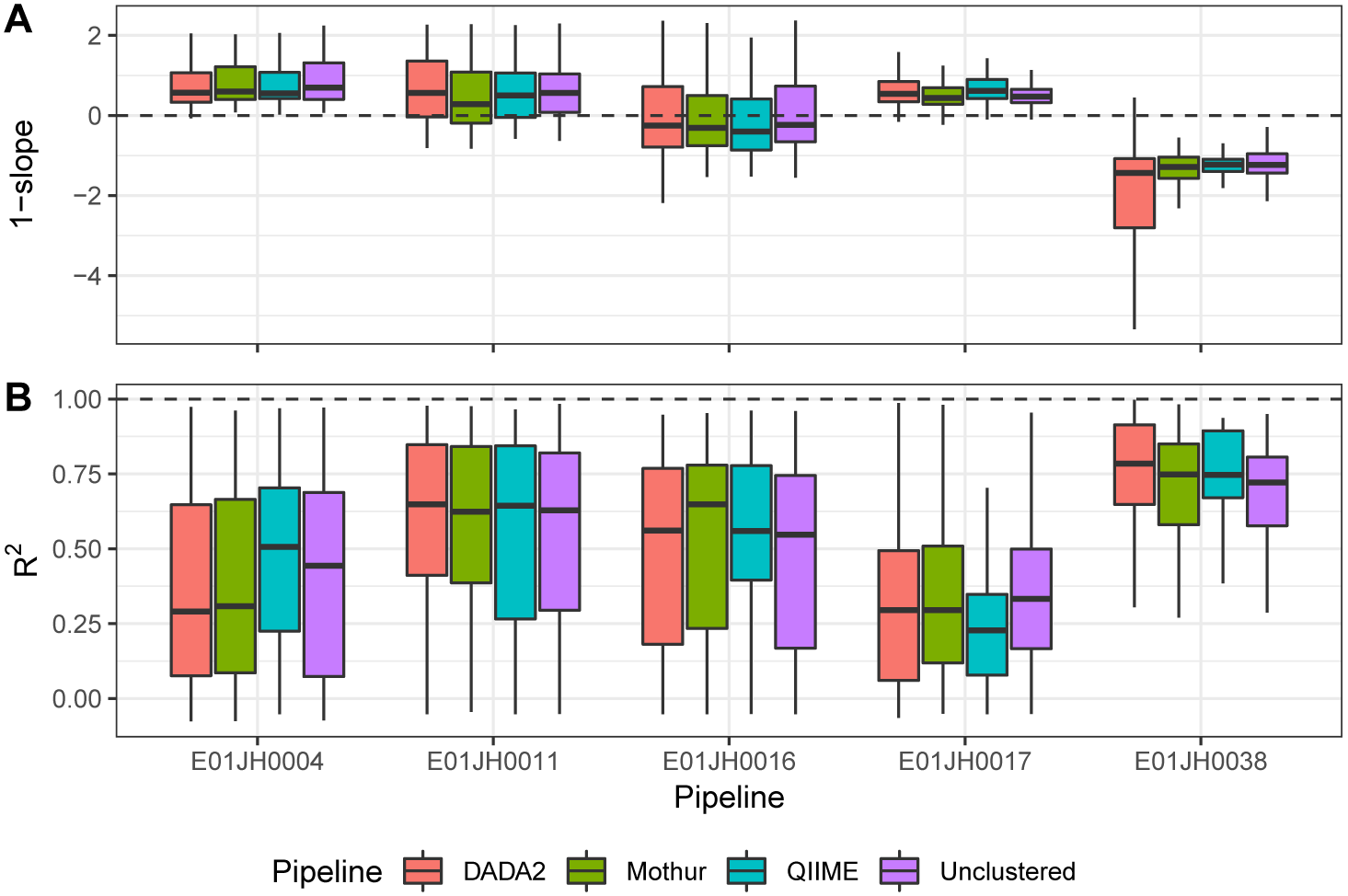
Feature-level log-fold change error bias (A) and variance (B) metric distribution by subject and pipeline. The bias (1*-slope*) and variance (*R*^2^) metrics are derived from the linear model fit to the estimated and expected log fold-change values for individual features. Boxplot outliers, 1.5*× IQR* from the median were excluded from the figure to prevent extreme metric values from obscuring metric value visual comparisons.

The log fold-change error distribution was consistent across pipelines (Fig. 10B). There was a long tail of high error features in the error distribution for all pipelines and individuals. The log fold-change estimates responsible for the long tail could not be attributed to specific titration comparisons. Additionally, we compared log-fold change error distributions for log-fold change estimates using different normalization methods. Error rate distributions, including the long tails, were consistent across normalization methods. Furthermore, as the long tail was observed for the unclustered data as well, the log-fold change estimates contributing to the long tail are likely due to a bias associated with the molecular laboratory portion of the measurement process and not the bioinformatic pipelines. Exploratory analysis of the relationship between the log fold-change estimates and expected values for individual features indicated that the long tails were attributed to feature specific performance.

Feature-level log fold-change bias and variance metrics were used to compare pipeline performance (Fig. 10). Similar to relative abundance feature-level bias and variance metrics are defined as the 1 *-slope* and *R*2 for linear models of the estimated and expected log fold-change for individual features and all titration comparisons. For the bias metric, 1 *-slope*, the desired value is 0 (i.e., log fold-change estimate = log fold-change expected), with negative values indicating the log-fold change was consistently underestimated and positive values consistently overestimated. The linear model *R*^2^ value was used to characterize the feature-level log fold-change variance as it indicates consistency between log fold-change estimates and expected values across titration comparisons. To compare bias and variance metrics across pipelines mixed-effects models were used. The log fold-change bias and variance metrics were not significantly different between pipelines (Bias: F = 0, 2.51, p = 0.99, 0.08, 10B, Variance: F = 47.39, 0.23, p = 0, 0.8, Fig. 10C). We also evaluated whether poor feature-level metrics could be attributed to specific clades for taxonomic groups. Similar to the relative abundance estimate, while a phylogenetic signal was detected for both the bias and variance metrics, no specific taxonomic groups or phylogenetic clades that performed poorly were identified.

## Discussion

We assessed the quantitative and qualitative characteristics of count tables generated using different bioinformatic pipelines and 16S rRNA marker-gene survey mixture dataset. The mixture dataset followed a two-sample titration mixture design, where DNA collected before and after exposure to pathogenic *Escherichia coli* from five vaccine trial participants (subjects) were mixed following a *log*_2_ dilution series (Fig. 1). Qualitative count table characteristics were assessed using relative abundance information for features observed only in titrations and unmixed samples. We quantitatively assed count tables by comparing feature relative and differential abundance to expected values.

### Count Table Assessment Demonstration

We demonstrated our novel assessment approach by evaluating count tables generated using different bioinformatic pipelines, QIIME, Mothur, and DADA2. The Mothur pipeline uses *de novo* clustering for feature inference [22, 23]. Pairwise distances used in clustering are calculated using a multiple sequence alignment. The quality filtered paired-end reads are merged into contigs. The pipeline then aligns contigs to a reference multiple sequence alignment and removes uninformative positions in the multiple sequence alignment. The QIIME pipeline uses open-reference clustering where merged paired-end reads are first assigned to reference cluster centers [24, 25]. Next QIIME clusters unassigned reads *de novo*. Unlike Mothur, the QIIME clustering method uses pairwise sequence distances calculated from pairwise sequence alignments. As a result, the QIIME pairwise distances are calculated using the full ˜436 bp sequences whereas Mothur pairwise distances were calculated using a 270 bp multiple sequence alignment. The DADA2 pipeline uses a probability model and maximization expectation algorithm for feature inference [7]. Unlike distance-based clustering methods employed by the Mothur and QIIME pipelines, DADA2 parameters determine if low abundance sequences are grouped with a higher abundance sequence. As a control, we compared our quantitative assessment results for the three pipelines to a count table of unclustered features. The unclustered features were generated using the Mothur pipeline preprocessing methods.

#### Quantitative Assessment

While the relative abundance bias metric was significantly different between pipelines overall, pipeline choice had minimal impact on the quantitative assessment results when accounting for subject-specific effects. Outlier features, those with extreme quantitative analysis bias and variance metrics, were observed for all pipelines and both relative and differential abundance assessments. Outlier features are not likely a pipeline artifact as they were observed in count tables generated using the unclustered pipeline as well as standard bioinformatic pipelines. We were unable to attribute outlier features to relative abundance values, log fold-change between unmixed samples, and sequence GC content. Features with extreme metric values were not limited to any specific taxonomic group or phylogenetic clade. Outlier features could not be attributed to bioinformatic pipelines and are likely due to biases in the molecular biology part of the measurement process. PCR amplification is a well-known bias in the molecular biology part of the measurement process. Mismatches in the primer binding regions impact PCR efficiency and are a potential cause for poor feature-specific performance [26]. Additional research is needed before outlier features are attributed to mismatches in the primer binding regions.

#### Qualitative Assessment

The qualitative assessment evaluated whether features only observed in unmixed samples or titrations could be explained by sampling alone. Features present only in titrations or unmixed samples not due to random sampling are bioinformatic pipeline artifacts. These artifacts can be categorized as false negative or false positive features. A false negative occurs when a lower abundance sequence representing an organism within the sample is clustered with a higher abundance sequence from a different organism. False positives are sequencing or PCR artifacts not appropriately filtered or assigned to an appropriate feature by the bioinformatic pipeline.

Count table sparsity, the proportion of zero-valued cells, provides additional insight into the qualitative assessment results. A high rate of false negative features is a potential explanation for DADA2 count table’s poor performance in the qualitative assessment and comparable sparsity to the other pipelines despite having significantly fewer features (Fig. 7 and Table 1). The DADA2 feature inference algorithm may be aggressively grouping lower abundance true sequences with higher abundance sequences. As a result, the low abundance sequences are not present in samples leading to increased sparsity and higher abundance unmixedand titrationspecific features. Adjusting the DADA2 parameters, specifically the OMEGA A parameter in setDadaOpt. Along these lines, the DADA2 documentation states that the default setting for OMEGA A is conservative to prevent false positives at the cost of increasing false negatives [7].

False positive features provide an explanation for Mothur and QIIME pipelines having lower proportion of unmixed- and titration-specific features not explained by sampling but high sparsity (Fig. 7 and Table 1). The statistical tests used to determine if the specific features could be explained by sampling alone only considers feature abundance. Therefore, the statistical test is not able to distinguish between true low abundance unmixed- and titration-specific features and low abundance sequence artifacts. Mothur and QIIME count tables have ten times and three times more features compared to DADA2, respectively (Table 1). While microbial abundance distributions are known to have long tails, it is likely that the observed sparsity is an artifact of the 16S rRNA sequencing measurement process. Similarly, significantly more features than expected are commonly observed for mock community benchmarking studies evaluating the QIIME and Mothur pipelines [20].

False positive features can be reduced, but not eliminated, using smaller amplicon and prevalence filtering. The 16S rRNA region sequenced in the study is larger than the region the *de-novo*, and open clustering pipelines were developed for, potentially explaining the higher than expected sparsity [20]. Kozich et al. [20] reduced the sequence error rate from 0.29% to 0.06% by using paired-end reads that completely overlap. The larger region used in this study has a smaller overlap between the forward and reverse reads. As a result, merging the forward and reverse reads did not allow for sequence error correction that occurs when a smaller amplicon is used. However, even when targeting smaller regions of the 16S rRNA gene both the *de-novo* (Mothur) and open-reference clustering (QIIME) pipelines produced count tables with significantly more features than expected in evaluation studies using mock communities. Prevalence filtering is used to exclude low abundance features, predominantly measurement artifacts [27]. For example, a study exploring the microbial ecology of the Red-necked stint *Calidris ruficollis*, a migratory shorebird, used a hard filter to validate their study conclusions are not biases by false positive features. The study authors compared results with and without prevalence filter ensuring that the study conclusions were not biased by using the arbitrary filter or including the low abundant features [28].

### Using Mixtures to Assess 16S rRNA Sequencing

Mixtures of environmental samples have previously been used to assess RNAseq and microarray gene expression measurements. However, this is the first time mixtures have been used to assess microbiome measurement methods. Using our mixture dataset we developed novel methods for assessing marker-gene-survey computational methods. Our quantitative assessment allowed for the characterization of relative abundance values using a dataset with a larger number of features and dynamic range compared to mock community assessments. As a result, we identified previously unknown feature specific biases. Based on our subject-specific results observation, we recommend that studies using stool samples seeking inferences in a longitudinal series of multiple subjects carefully estimate bacterial DNA proportions and adjust inferences accordingly. Additionally, our qualitative assessment results, when combined with sparsity information provide a new method for evaluating how well bioinformatic pipelines account for sequencing artifacts without loss of true biological sequences.

There were also limitations using our mixture dataset. These limitations included: Lack of agreement between the proportion of unmixed samples titrations and the mixture design. The number of features used in the different analysis. These limitations are described below along with recommendations for addressing them in future studies.

Differences in the proportion of prokaryotic DNA in the samples used to generate the two-sample titrations series resulted in differences between the true mixture proportions and mixture design. We attempted to account for differences in mixture proportion from mixture design by estimating mixture proportions using sequence data. Similar to how the proportion of mRNA in RNA samples was used in a previous mixture study [15]. We used an assay targeting the 16S rRNA gene to detect changes in the concentration of prokaryotic DNA across titrations, but were unable to quantify the proportion of prokaryotic DNA in the unmixed samples using qPCR data. Using the 16S sequencing data we inferred the proportion of prokaryotic DNA from the POST sample in each titration. However, the uncertainty and accuracy of the inference method are not known resulting in an unaccounted for error source.

A better method for quantifying sample prokaryotic DNA proportion or using samples with consistent proportions would increase confidence in the expected value and in-turn error metric accuracy. Limitations in the prokaryotic DNA qPCR assay’s concentration precision limits the assay’ssuitability for use in mixture studies. Digital PCR provides a more precise alternative to qPCR and is, therefore, a more appropriate method. Alternatively using samples where the majority of the DNA is prokaryotic would minimize this issue. Mixtures of environmental samples can also be used to assess shotgun metagenomic methods as well. As shotgun metagenomics is not a targeted approach, differences in the proportion of prokaryotic DNA in a sample would not impact the assessment results in the same way as 16S rRNA marker-gene-surveys.

Using samples from a vaccine trial allowed for the use of a specific marker with an expected response, *E. coli*, during methods development. However, the high level of similarity between the unmixed samples resulted in a limited number of features that could be used in the quantitative assessment results. Using more diverse samples to generate mixtures would address this issue.

## Conclusions

Our two-sample-titration dataset and assessment methods can be used to evaluate and characterize bioinformatic pipelines and clustering methods. The sequence dataset presented in this study can be processed with any 16S rRNA bioinformatic pipeline. Our quantitative and qualitative assessment can then be performed on the count table and the results compared to those obtained using the pipelines presented here. The three pipelines we evaluated produced sets of features varying in total feature abundance, number of features per samples, and total features. The objective of any pipeline is to differentiate true biological sequences from measurement process artifacts. In general, based on our evaluation results we suggest using DADA2 for feature-level abundance analysis, e.g. differential abundance testing. While DADA2 performed poorly in our qualitative assessment, the pipeline performed better in the quantitative assessment compared to the other pipelines. Additionally, the DADA2 poor qualitative assessment results due to false-negative features are unlikely to negatively impact feature-level abundance analysis. When determining which pipeline to use for a study, users should consider whether minimizing false positives (DADA2) or false negatives (Mothur) is more appropriate for their study objectives. When a sequencing dataset is processed using DADA2, the user can be more confident that an observed feature represents a member of the microbial community and not a measurement artifact. Pipeline parameter optimization could address DADA2 false-negative issue. For the Mothur and QIIME pipelines, prevalence filtering will reduce the number of false-positive features. Feature-level results for any 16S rRNA marker-gene survey should be interpreted with care, as the biases responsible for poor quantitative assessment are unknown. Addressing both of these issues requires advances in both the molecular biology and computational components of the measurement process.

## Methods

### Titration Validation

qPCR was used to validate volumetric mixing and check for differences in the proportion of prokaryotic DNA across titrations. To ensure the two-sample titrations were volumetrically mixed according to the mixture design, independent ERCC plasmids were spiked into the unmixed PRE and POST samples [29] (NIST SRM SRM 2374) (Table 2). The ERCC plasmids were resuspended in 100 *µL* tris-EDTA buffer and 2 *µL* of resuspended plasmids was spiked into the appropriate unmixed sample. Plasmids were spiked into unmixed samples after unmixed sample concentration was normalized to 12.5 *ng/µL*. POST sample ERCC plasmid abundance was quantified using TaqMan gene expression assays (FAM-MGB, Catalog # 4448892, ThermoFisher) specific to each ERCC plasmid and TaqMan Universal MasterMix II (Catalog # 4440040, ThermoFisher Waltham, MA USA).

To check for differences in the proportion of bacterial DNA in the PRE and POST samples, titration bacterial DNA concentration was quantified using the Femto Bacterial DNA quantification kit (Zymo Research, Irvine CA). All samples were run in triplicate along with an in-house *E. coli* DNA *log*_10_ dilution standard curve. qPCR assays were performed using the QuantStudio Real-Time qPCR (ThermoFisher). Amplification data and Ct values were exported as tsv files using QuantStudio Design and Analysis Software v1.4.1. Statistical analysis was performed on the exported data using custom scripts in R [30]. The qPCR data and scripts used to analyze the data are available at https://github.com/nate-d-olson/mgtst_pub.

### Sequencing

The 45 samples (seven titrations and two unmixed samples for each of five subjects) were processed using the Illumina 16S library protocol (16S Metagenomic Sequencing Library Preparation, posted date 11/27/2013, downloaded from https://support.illumina.com). This protocol specifies an initial 16S rRNA gene PCR, followed by a sample indexing PCR, normalization, and sequencing.

A total of 192 16S rRNA PCR assays were run including four replicates per sample and 12 no-template controls, using Kapa HiFi HotStart ReadyMix reagents (KAPA Biosystems, Inc. Wilmington, MA). The initial PCR assay targeted the V3-V5 region of the 16S rRNA gene, Bakt 341F and Bakt 806R [10]. The V3-V5 region is 464 base pairs (bp) long, with forward and reverse reads overlapping by 136 bp, using 2 X 300 bp paired-end sequencing [31] (http://probebase.csb.univie.ac.at). Primer sequences include overhang adapter sequences for library preparation (forward primer 5’-TCG TCG GCA GCG TCA GAT GTG TAT AAG AGA CAG CCT ACG GGN GGC WGC AG 3-’ and reverse primer 5’-GTC TCG TGG GCT CGG AGA TGT GTA TAA GAG ACA GGA CTA CHV GGG TAT CTA ATC C -3’). For quality control, the PCR product was verified using agarose gel electrophoresis to check amplicon size. Concentration measurements were made after the initial 16S rRNA PCR, the indexing PCR, and normalization steps. DNA concentration was measured using the QuantIT Picogreen dsDNA Kit (Cat # P7589, ThermoFisher Scientific) and fluorescent measurements were made with a Synergy2 Multi-Detection MicroPlate Reader (BioTek Instruments, Inc, Winooski, VT).

Initial PCR products were purified using 0.8X AMPure XP beads (Beckman Coulter Genomics, Danvers, MA) following the manufacturer’s protocol. After purification, the 192 samples were indexed using the Illumina Nextera XT index kits A and D (Illumina Inc., San Diego CA) and then purified using 1.12X AMPure XP beads. Prior to pooling purified sample concentration was normalized using SequalPrep Normalization Plate Kit (Catalog n. A10510-01, Invitrogen Corp., Carlsbad, CA), according to the manufacturer’s protocol. Pooled library concentration was checked using the Qubit dsDNA HS Assay Kit (Part# Q32851, Lot# 1735902, ThermoFisher, Waltham, MA USA). Due to the low pooled amplicon library DNA concentration, a modified protocol for low concentration libraries was used. The library was run on an Illumina MiSeq, and base calls were made using Illumina Real Time Analysis Software version 1.18.54. Sequencing data quality control metrics for the 384 fastq sequence files (192 samples with forward and reverse reads) were computed using the Bioconductor Rqc package [32, 33]. The sequence data was deposited in the NCBI SRA archive under Bioproject PRJNA480312. Individual SRA run accession numbers and metadata in Supplemental Table.

### Sequence Processing

Sequence data were processed using four bioinformatic pipelines: a *de-novo* clustering method Mothur [23], an open-reference clustering method QIIME [25], and a sequence inference method DADA2 [7], and unclustered sequences as a control. The code used to run the bioinformatic pipelines is available at https://github.com/nate-d-olson/mgtst_pipelines.

The Mothur pipeline follows the developer’s MiSeq SOP [23, 20]. The pipeline was run using Mothur version 1.37 (http://www.mothur.org/). We sequenced a larger 16S rRNA region, with smaller overlap between the forward and reverse reads, than the 16S rRNA region the SOP was designed. Pipeline parameters modified to account for difference in overlap are noted for individual steps below. The Makefile and scripts used to run the Mothur pipeline are available https://github.com/nate-d-olson/mgtst_pipelines/blob/master/code/mothur. The Mothur pipeline includes an initial preprocessing step where the forward and reverse reads are trimmed and filtered using base quality scores and were merged into single contigs for each read pair. The following parameters were used for the initial contig filtering, no ambiguous bases, max contig length of 500 bp, and max homopolymer length of 8 bases. For the initial read filtering and merging step, low-quality reads were identified and filtered from the dataset based on the presence of ambiguous bases, failure to align to the SILVA reference database (V119, https://www.arb-silva.de/) [34], and identification as chimeras. Prior to alignment, the SILVA reference multiple sequence alignment was trimmed to the V3-V5 region, positions 6,388 and 25,316. Chimera filtering was performed using Uchime (version v4.2.40) without a reference database [35]. OTU clustering was performed using the OptiClust algorithm with a clustering threshold of 0.97 [22]. The RDP classifier implemented in Mothur was used for taxonomic classification against the Mothur provided version of the RDP v9 training set [36].

The QIIME open-reference clustering pipeline for paired-end Illumina data was performed according to the online tutorial (Illumina Overview Tutorial (an IPython Notebook): open reference OTU picking and core diversity analyses, http://qiime.org/tutorials/) using QIIME version 1.9.1 [25]. Briefly, the QIIME pipeline uses fastq-join (version 1.3.1) to merge paired-end reads [37] and the Usearch algorithm [38] with Greengenes database version 13.8 with a 97% similarity threshold [39] was used for open-reference clustering.

DADA2, an R native pipeline was also used to process the sequencing data [7]. The pipeline includes a sequence inference step and taxonomic classification using the DADA2 implementation of the RDP naÏve Bayesian classifier [36] and the SILVA database V123 provided by the DADA2 developers [34, https://benjjneb.github.io/dada2/training.html].

The unclustered pipeline was based on the Mothur *de-novo* clustering pipeline, where the paired-end reads were merged, filtered, and then dereplicated. Reads were aligned to the reference Silva alignment (V119, https://www.arb-silva.de/), and reads failing alignment were excluded from the dataset. Taxonomic classification of the unclustered sequences was performed using the same RDP classifier implemented in Mothur used for the *de-novo* pipeline. To limit the size of the dataset the most abundant 40,000 OTUs (comparable to the Mothur dataset), across all samples, were used as the unclustered dataset.

### Titration Proportion Estimates

The following linear model (2) was used to infer the proportion of prokaryotic DNA, *θ*, in each titration. Where **Q**_*i*_ is a vector of titration *i* feature relative abundance estimates and **Q**_*pre*_ and **Q**_*post*_ are vectors of feature relative abundance estimates for the unmixed PRE and POST samples. Feature relative abundance estimates were calculated using a negative binomial model.

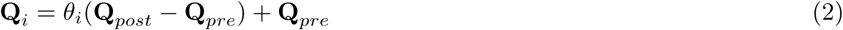

To fit the model and prevent uninformative and low abundance features from biasing *θ* estimates, only features meeting the following criteria were used. Features included in the model were observed in at least 14 of the 28 total titration PCR replicates (4 replicates per 7 titrations), demonstrated greater than 2-fold difference in relative abundance between the PRE and POST samples, and were present in either all four or none of the PRE and POST PCR replicates.

16S rRNA sequencing count data is known to have a non-normal mean-variance relationship resulting in poor model fit for standard linear regression [14]. Generalized linear models provide an alternative to standard least-squares regression. The above model is additive and therefore *θ*_*i*_ cannot be directly inferred in log-space. To address this issue, we fit the model using a standard least-squares regression then obtained non-parametric 95 % confidence intervals for the *θ* estimates by bootstrapping with 1000 replicates.

### Qualitative Assessment

Our qualitative measurement assessment evaluated features only observed in unmixed samples (PRE or POST), *unmixed-specific*, or titrations, *titration-specific*. *Unmixed*or *titration-specific* features are due to differences in sampling depth (number of sequences) between the unmixed samples and titrations, artifacts of the feature inference process, or PCR/sequencing artifacts. Measurement process artifacts should be considered false positives or negatives. Hypothesis tests were used to determine if differences in sampling depth could account for *unmixed-specific* and *titration-specific* features. p-values were adjusted for multiple comparisons using the Benjamini & Hochberg method [40]. For *unmixed-specific* features, the binomial test was used to evaluate if true feature relative abundance is less than the expected relative abundance. A binomial test could not be used to evaluate *titration-specific* features, as the hypothesis would be formulated as such. Given observed counts and the titration total feature abundance, the true feature relative abundance is equal to 0. As non-zero counts were observed the true feature proportion is non-zero, and the test always fails. Therefore, we formulated a Bayesian hypothesis test for *titration-specific* features.

A Bayesian hypothesis test was used to evaluate if the true feature proportion is less than the minimum detected proportion. The Bayesian hypothesis test was formulated using equation (3). Which when assuming equal priors, *P* (*π < π*_*min*_) = *P* (*π ≥ π*_*min*_), reduces to (4). For equations (3) and (4) *π* is the true feature proportion, *π*_*min*_ is the minimum detected proportion, *C* is the expected feature counts, and *C*_*obs*_ is the observed feature counts. Simulation was used to generate possible values of *C*, assuming *C* has a binomial distribution given the observed sample total feature abundance, and a uniform probability distribution for *π* between 0 and 1. *π*_*min*_ was calculated using the mixture equation (1) where *q*_*pre,j*_ and *q*_*post,j*_ are *min*(**Q**_*pre*_) and *min*(**Q**_*post*_) across all features for a subject and pipeline. Our assumption is that *π* is less than *π*_*min*_ for features not observed in unmixed samples due to random sampling.

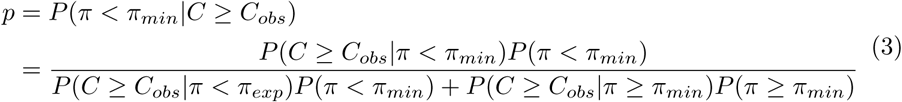

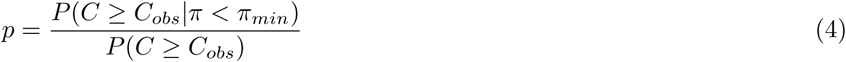

### Quantitative Assessment

For quantitative assessment, we compared observed relative abundance and log fold-changes to expected values derived from the titration experimental design. Feature average relative abundance across PCR replicates was calculated using a negative binomial model, and used as observed relative abundance values (*obs*) for the relative abundance assessment. Average relative abundance values were used to reduce PCR replicate outliers from biasing the assessment results. Equation (1) and inferred *θ* values were used to calculate the expected relative abundance values (*exp*). Relative abundance error rate is defined as *|exp - obs|/exp*.

We developed bias and variance metrics to assess feature performance. The feature-level bias and variance metrics were defined as the median error rate and robust coefficient of variation (*RCOV* = *IQR/median*) respectively. Mixed-effects models were used to compare feature-level error rate bias and variance metrics across pipelines with subject as a random effect. Extreme feature-level error rate bias and variance metric outliers were observed, these outliers were excluded from the mixed effects model to minimize biases due to poor model fit and were characterized independently.

Log fold-change between samples in the titration series including PRE and POST were compared to the expected log fold-change values to assess differential abundance log fold-change estimates. Log fold-change estimates were calculated using EdgeR [41, 42]. Expected log fold-change for feature *j* between titrations *l* and *m* is calculated using equation (5), where *θ* is the proportion of POST bacterial DNA in a titration, and *q* is feature relative abundance. For features only present in PRE samples the expected log fold-change is independent of the observed counts for the unmixed samples and is calculated using (6). Features only observed in POST samples, *POST-specific*, expected log fold-change values can be calculated in a similar manner. However, *POST-specific* features were rarely observed in more than one titration and therefore were not suitable for use in our assessment. Due to a limited number of *PRE-specific* features, both *PRE-specific* and *PRE-dominant* features were used in the differential abundance assessment. *PRE-specific* features were defined as features observed in all four PRE PCR replicates and not observed in any of the POST PCR replicates and *PRE-dominant* features were also observed in all four PRE PCR replicates and observed in one or more of the POST PCR replicates with a log fold-change between PRE and POST samples greater than 5.

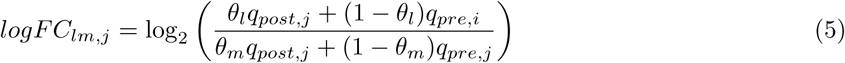

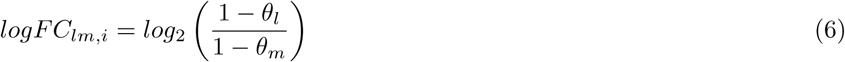

## Declarations

### Ethics approval and consent to participate

Not applicable.

### Consent for publication

Not applicable.

### Availability of data and material

Sequence data was deposited in the NCBI SRA archive under Bioproject PRJNA480312. Individual SRA run accession numbers and metadata in Supplemental Table. The code used to run the bioinformatic pipelines is available at https://github.com/nate-d-olson/mgtst_pipelines. Scripts used to analyze the data are available at https://github.com/nate-d-olson/mgtst_pub.

### Authors’ contributions

NDO, HCB, OCS, MS, and WT designed the experiment, SL and SH performed the laboratory work. NDO, HCB and MS analyzed the data. NDO and HCB wrote the manuscript. All authors provided feedback on manuscript drafts and approved the final manuscript.

## Acknowledgements

The authors would like to thank Mihai Pop, Scott Pine, Scott Jackson, Justin Zook, and Prachi Kulkarni for feedback on manuscript drafts. Opinions expressed in this paper are the authors and do not necessarily reflect the policies and views of NIST, or affiliated venues. Certain commercial equipment, instruments, or materials are identified in this paper in order to specify the experimental procedure adequately. Such identification is not intended to imply recommendations or endorsement by NIST, nor is it intended to imply that the materials or equipment identified are necessarily the best available for the purpose. Official contribution of NIST; not subject to copyrights in USA.

